# Comparative genomics: Dominant coral-bacterium *Endozoicomonas acroporae* metabolizes dimethylsulfoniopropionate (DMSP)

**DOI:** 10.1101/519546

**Authors:** Kshitij Tandon, Pei-Wen Chiang, Chih-Ying Lu, Naohisa Wada, Shan-Hua Yang, Ya-Fan Chan, Ping-Yun Chen, Hsiao-Yu Chang, Ming-Shean Chou, Wen-Ming Chen, Sen-Lin Tang

## Abstract

Dominant coral-associated *Endozoicomonas* bacteria species are hypothesized to play a role in the coral-sulfur cycle by metabolizing Dimethylsulfoniopropionate (DMSP) into Dimethylsulfide (DMS); however, no sequenced genome to date harbors genes for this process. In this study, we assembled high-quality (>95% complete) genomes of strains of a recently added species *Endozoicomonas acroporae* (Acr-14^T^, Acr-1 and Acr-5) isolated from the coral *Acropora muricata* and performed comparative genomic analysis on genus *Endozoicomonas*. We identified the first DMSP CoA-transferase/lyase—a *dddD* gene homolog found in all *E. acroporae* strains—and functionally characterized bacteria capable of metabolizing DMSP into DMS via the DddD cleavage pathway using RT-qPCR and gas chromatography (GC). Furthermore, we demonstrated that *E. acroporae* strains can use DMSP as the sole carbon source and have genes arranged in an operon-like manner to link DMSP metabolism to the central carbon cycle. This study confirms the role of *Endozoicomonas* in the coral sulfur cycle.

## Introduction

Coral reefs are one of the most diverse ecosystems on Earth, with over 800 different coral species known to date. Of these, corals in the genus *Acropora* are some of the most abundant reef-building corals across the Indo-Pacific region [1]. These corals are also some of the significant producers of dimethylsulfoniopropionate (DMSP) [2, 3]. DMSP is an organic sulfur compound present in significant amounts in animals that harbor symbiotic algae such as scleratinian corals and giant clams [2]. DMSP is present in coral tissues, mucus and endosymbiotic dinoflagellates (Symbiodiniaceae) [4, 5]. In marine algae, DMSP acts as a protectant from various stresses such as oxidative and osmotic stress [6]. Moreover, DMSP also acts as an attractant for specific bacterial groups that have been reported to be part of coral-associated bacterial communities and underpin coral health [7].

Once released from marine planktonic dinoflagellates, most of the DMSP is emanated to surrounding water where it is readily available for microbial catabolic conversion as a source of reduced carbon and sulfur [8, 9]. DMSP is a central molecule for marine sulfur cycling and is degraded by bacteria via two pathways, a cleavage pathway and a demethylation pathway [9, 10]. The majority (∼75%) of the DMSP is metabolized via the demethylation/demethiolation pathway, producing methylmercaptopropionate [11, 12, 13]. The cleavage pathway, which accounts for the remaining ∼25%, produces dimethylsulfide (DMS)—a climate-active gas and acrylic acid [13, 14, 15]. Genes and an enzyme (DddL) involved in the cleavage pathway has been identified in *Sulfitobacter* [16]. Another gene, *dddD*, has been identified in *Marinomonas* sp., which cleaves DMSP to produce DMS and 3-hydroxypropionate (3HP), not the usual acrylate [13]. Moreover, DMSP metabolism yielding DMS and acrylate was recently also identified in coccolithophore algae [17]. The DMS produced after cleavage of DMSP is then released in the surrounding water [12].

Coral-associated bacterial communities are highly abundant, dynamic and diverse [18, 19, 20, 21]. These bacterial communities can be found in various niches associated with corals like coral mucus [22], spaces in the skeleton [23, 24, 25], and within coral tissues [26, 27, 28]. Raina et al (2008) [8] confirmed that coral-associated bacteria have the potential to metabolize organic sulfur compounds present in the coral tissues. They inferred that the majority of the DMSP-degrading bacteria belongs to class *Gammaproteobacteria*, including *Alteromonas*-, *Arhodomonas-, Idiomarina-, Pseudomonas-*, and *Spongiobacter (Endozoicomonas)*-related organisms. Of these, *Arhodomonas-, Pseudomonas-*, and *Roseobacter-*related species harbor a DMSP lyase—i.e. the *dddD* gene.

Of all the organisms in the tremendously diverse coral holobionts, species in the genus *Endozoicomonas* (class: *Gammaproteobacteria*) have been studied widely for their genomics and functional and ecological roles [29, 30]. *Endozoicomonas* species were found to be abundant in the coral holobionts across the globe [31] and have been hypothesized to be potential indicators of coral health [32]—they have high abundance in healthy corals and a relatively low abundance in diseased or stressed corals [33, 34, 35]. Furthermore, nearly complete and draft genomes of *Endozoicomonas* species isolated from corals and other marine invertebrates [29, 34, 37] have been recently assembled. First, a nearly complete genome of the *Endozoicomonas* isolate *Endozoicomonas montiporae* CL-33^T^ from the encrusting pore coral *Montiporae aequituberculata* has provided insights into how this bacterium can interact with its host from outside to inside of the coral cell with the help of potential effector proteins [30]. A recent comparative genomics analysis also identified a high number of Type III secretion system-related genes and that most gene ontology terms were associated with the generic transport of molecules and those genomes of *Endozoicomonas* species show high plasticity [30]. *Endozoicomonas* species have been hypothesized to play a role in the coral sulfur cycle by effectively metabolizing DMSP into DMS [38, 39]. However, no study has confirmed the genus’ role and no sequenced genome has harbored any genes in this process [30]. Hence, the role of this coral endosymbiont in the coral sulfur cycle remains elusive.

In this study, we assembled high quality and nearly complete draft genomes of newly added *Endozoicomonas acroporae* strains and profiled its abundance in different coral species in the Indo-pacific region. We identified and functionally characterized the first DMSP Co-A transferase/lyase; dddD homolog in all *E. acroporae* strains. Furthermore, we provide conclusive evidence that *E. acroporae* has a role in the coral sulfur cycle by effectively metabolizing DMSP into DMS and is able to use DMSP as the sole carbon source for growth and survival.

## Materials and Methods

### Culturing and whole-genome sequencing of *E. acroporae* Acr-1, Acr-5 and Acr-14^T^

Strains of *E. acroporae* Acr-1, Acr-5 and Acr-14^T^ were isolated from the coral *Acropora muricata* from Kenting, off the southern coast of Taiwan, and cultured using a method described previously [40]. Genomic DNA isolation was performed using the cetyltrimethylammonium bromide (CTAB) method [41]; the quality of the isolated DNA was assessed using NanoDrop 1000 (Thermo Scientific, USA). High quality DNA was sent to the core sequencing facility at Biodiversity Research Center, Academia Sinica (BRCAS), Taiwan for whole-genome sequencing on the Illumina Miseq platform, with a TRUSeq DNA paired-end library generated to achieve an insert size of 500 bp.

### Genome assembly and annotation

Reads obtained from Illumina MiSeq were quality-filtered and trimmed (Phred score ≥ 30) using NGS QC toolkit v2.3.3 [42]. Quality-filtered and trimmed reads were *de novo* assembled using CLC Genomics Workbench version 1.10.1 (Qiagen) with a bubble size of 40 and automatic word size enabled. Minimum contig length was set to ≥500 bp (no scaffolding was performed). Assembled genomes were quality checked for completeness, contamination, and heterogeneity using CheckM [43]. Other *Endozoicomonas* species genomes were downloaded from the NCBI genomes database (last accessed January 2018). Gene prediction was performed using Prokka [44] on all genomes used in this study to avoid bias from different gene prediction methods. Predicted genes were annotated by rapid annotation using the Subsystem Technology (RAST) Server [45] with gene calls from Prokka preserved.

### Identification of genomic characteristics

The *in-silico* DNA-DNA hybridization (DDH) percentages among the genomes of genus *Endozoicomonas* were calculated using the genome-to-genome distance calculator from the DSMZ server [46], and the Average Nucleotide Identity (ANI) (https://enveomics.ce.gatech.edu/ani/) calculator was used to calculate the ANI values. Further, Amino-Acid Identity (AAI) among the genomes was calculated with CompareM (https://github.com/dparks1134/CompareM). Clustered Regularly Interspaced Short Palindromic Repeat (CRISPR) structures in all genomes were identified using Prokka; prophages and phages within the genomes were identified using the PHAge Search Tool (PHAST) [47], which classified the phages as intact, incomplete and questionable. Type III secretion system (T3SS) proteins were identified by EffectiveT3 [48] using EffectiveDB with the animal classification module and selective (0.9999) restriction value method enabled. Insertion Sequence (IS) elements in the genomes of *E. acroporae* strains and *E. montiporae* were identified using the ISfinder database (https://www.is.biotoul.fr) with blastn and an e-value threshold of 1e-5.

### 16S rRNA gene phylogenetic analysis

To determine robust phylogenetic relationships within the genus *Endozoicomonas*, all available 16S rRNA sequences (80 in number) were downloaded from the NCBI taxonomy database, for which host information was available to understand the distribution of *Endozoicomonas* species in different marine invertebrates and identify the position of *E. acroporae* strains within the genus *Endozoicomonas*. Sequences were aligned using MUSCLE [49] in MEGA7 [50] with default parameters, and conserved regions were extracted manually. Furthermore, the conserved regions were again aligned using the above method. The phylogenetic tree was constructed using the maximum-likelihood method with Kimura 2-parameter and Gamma distributed with invariant sites enabled in MEGA7 with 1000 bootstraps.

### *E. acroporae* distribution and abundance in different coral species from the Indo-Pacific region

We analyzed microbial community data publically available from three different studies—1) our laboratory’s previous study [34]; 2) a study of coral-associated bacterial communities in the Red Sea [51]; and 3) coral-associated bacterial community from reefs in the east and west coast of Australia, including Ningaloo Reef, Lizard Island, reefs from the northern sector of the Great Barrier Reef, and Lorde Howe Island [52]—to profile the abundance of *E. acroporae* strains in different corals species from Penghu Archipelago, Taiwan, the Red Sea, Saudi Arabia, and east and west Australia. Operational Taxonomic Unit (OTU) abundance profiles and their representative sequences were obtained from supplementary materials in Shiu JH et al, 2017 [34], Ziegler M et al, 2016 [51], and Pollock FJ et al, 2018 [52]. We performed similarity searches on all the OTU sequences in the three studies against an in-house 16S rRNA gene sequence database of *Endozoicomonas* species with standalone blastn [53] and profiled the relative abundance of *E. acroporae* strains at the three locations with e-value <1e-5 and identity threshold ≥ 97%.

### Comparative genomics: Pan-genome analysis and core-genome phylogeny

Pan-genome analysis was performed with Roary [54] using GFF files of all the genomes obtained with Prokka. Core genome identification and alignment was performed on all genomes using the parameters *-i 80, -e, –n*. A core genome-based phylogenetic tree was constructed using Fasttree [55] with –nt, -gtr parameters and visualized in the GENETYX-Tree Program (https://www.genetyx.co.jp/).

### Identification of stress response genes, *dddD CoA-transferase/lyase*, and DMSP metabolism related operons

The Stress Response Subsystem was analyzed for distribution of different categories of stress responsive genes present in the genomes of *Endozoicomonas*. The Sulfur Metabolism Subsystem in the RAST analysis annotated a *dddD* gene capable of metabolizing DMSP into DMS within the “Sulfur Metabolism-no subcategory.” Furthermore, the presence of domains in the *dddD* gene and DMSP metabolism-related operon genes was determined with web-based Conserved Domain Search Service [56,57], NCBI, with default parameters.

### DMSP degradation by strain Acr-14^T^

*Endozoicomonas acroporae* Acr-14^T^ was cultivated on the Modified Marine Broth Version 4 (MMBV4) medium [29] with several modifications (Supplementary Table S1) at 25 °C for 48 hours. Carbon sources, 0.1 % maltose, and 0.5 mM DMSP were added to the enrichment culture of strain Acr-14^T^and kept at 25 °C for 24 hours (OD_600_ of ∼1.0 after incubation). The enrichment culture was centrifuged at 2000 × *g* for 10 minutes, and the supernatant was discarded. The enrichment culture pellet was washed with 1 ml fresh minimal medium (Supplementary Table S2) twice to remove MMBV4 medium containing DMSP. Two experimental groups were made for the test: (A) 1 ml of washed bacteria was resuspended in a 40 ml minimal medium containing 0.2% casamino acid and 1 mM DMSP; (B) 1 ml of washed bacteria was resuspended in a 40 ml minimal medium containing 0.2% casamino acid without DMSP. The cultures were sampled at 0, 16, 24 and 48 hours for RT-qPCR.

### Gene expression of *dddD* by RT-qPCR

Total RNA was extracted using a TRI-reagent solution (Invitrogen, Carlsbad, CA, USA). The culture samples (1-2 ml) were centrifuged for 1 min at 12000 × *g* and 4 °C following the manufacturer’s guidelines. The RNA pellet was air-dried and resuspended in nuclease-free water. Residual DNA was removed using a TURBO DNA-free Kit (Invitrogen). RNA quality was determined using NanoDrop ND-1000 UV-Vis Spectrophotometer (Nano-Drop Technologies). RNA integrity was assessed by electrophoresis on 1% agarose-guanidine thiocyanate gel. Complementary DNA (cDNA) was synthesized from purified RNA using the SuperScript IV First Strand Synthesis System for RT-qPCR (Invitrogen) following the manufacturer’s guidelines. For cDNA, RT and non-RT samples were screened for residual DNA contamination using the hypervariable V6V8 region of the bacterial 16S rRNA gene (U968F and U1391R).

A primer pair was made for the *dddD* gene based on the genome data of strain Acr-14^T^ for Quantitative PCR using DNASTAR Lasergene [58] and Primer-BLAST tool on BLAST search (NCBI): *dddD*-F (5’-ACCGCATCGCACCACTCAGG-3’) and *dddD*-R (5’-GGCCCCGGTTGTTTCATCAT-3’). The endogenous control was performed with *rpoD* gene, *rpoD*-F (5’-AAGGCGGTGGACAAGTTCG-3’) and *rpoD*-R (5’-GATGGTGCGGGCCTGGTCTG-3’). RT-qPCR assays were carried out using Applied Biosystems QuantStudio™5 Real-Time PCR System. The standard cycling program consisted of cycles of UDG activation at 50 °C for 2 minutes, initial denaturation-activation at 95 °C for 2 minutes and 40 cycles of denaturation at 95 °C for 10 seconds and annealing at 60 °C for 40 seconds using PowerUp SYBR Green Master Mix (Thermo Fisher Scientific, USA). A dissociation step was performed to confirm the specificity of the product and avoid the production of primer dimers. For all reactions, 10 ng of template DNA was added to a reaction of 10 μl. The 10 μl reactions contained 5 μl of PowerUp SYBR Green Master Mix (Thermo Fisher Scientific, USA), 3.4 μl of sterilized nuclease-free water, 0.3 μl each of the forward and reverse primers (final conc. 0.3 µM), and 1μl of DNA template. Each sample was performed in duplicate. The relative quantification of the expression ratio was calculated by the comparative 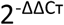 method. Differences between treatments were statistically tested using *t-test*.

### Quantification of released DMS

To assay the functional activity of the DddD protein in *Endozoicomonas acroporae* strain Acr-14^T^, cultures were first grown in MMB medium with 0.5 mM DMSP for 24 hours. After that, 3 ml culture was collected, spun down, and washed twice with minimal medium and then resuspended with 1 ml minimal medium. This 1 ml culture was then injected into sterile 60 ml vials sealed with a rubber stoppers containing designated medium Treatment (a): 20 ml minimal medium with 0.2 % casamino acid, and 1 mM DMSP; Treatment (c): 20 ml minimal medium with 0.2 % casamino acid, acting as the negative control; and a culture-free Treatment (b): 20 ml minimal medium with 0.2 % casamino acid, and 1 mM DMSP acted as the control. Vials were then incubated at 25 °C, 200 rpm, in dark. After incubation for 0, 24 and 48 hours, 1 ml of the head space air sample was collected and injected into a gas chromatograph (Shimadzu, GC-14B) fitted with a flame ionization detector (150 °C) and column (SGE, 60m × 0.53mm ID BP624 × 3.0 µm) to detect the DMS concentration.

### *E. acroporae* growth on DMSP as the sole carbon source

*E. acroporae* Acr-1, Acr-5, and Acr-14^T^ and *E. montiporae* CL-33^T^ strains were cultivated on MMB medium (along with 0.1% maltose for *E. montiporae* CL-33^T^) at 25 °C for 72 hours. The carbon source, 0.1 mM DMSP, was added to the enrichment culture of *E. acroporae* strains and kept at 25 °C for 24 hours (OD_600_ of ∼0.3 after incubation). 0.1 mM DMSP and 0.1% maltose were added to the enrichment culture of *E. montiporae* and kept at 25 °C for 24 hours (OD_600_ of ∼0.8 after incubation). The enrichment culture was centrifuged at 2000 × g for 10 minutes, and the supernatant was discarded. The enrichment culture pellet was washed twice by 1 ml fresh minimal medium. After resuspending with minimal medium, the enrichment cultures were added to the treatments and adjusted to OD_600_ of 0.06. All the treatments were kept at 25 °C, 200 rpm, and the OD_600_ of each treatment was recorded after incubating for 24, 48, and 72 hours.

*E. acroporae* Acr-14^T^, Acr-1, Acr-5, and *E. montiporae* CL-33^T^ were cultivated in minimal medium with 0.2% casamino acid and 0.1 mM DMSP to test their ability in usage DMSP as the sole carbon source. Treatments with different concentrations of DMSP were set to confirm whether *E. montiporae* CL-33^T^ can grow with DMSP or not. In these treatments, *E. montiporae* CL-33^T^ was cultivated in minimal medium with 0.2% casamino acid and 3 mM DMSP; 0.2% casamino acid and 7 mM DMSP; 0.2% casamino acid and 1% maltose as a positive control.

## Data Availability

Draft genomes of *E. acroporae* Acr-1 and *E. acroporae* Acr-5 have been deposited into GenBank under accession IDs SAUT00000000 and SAUU00000000, respectively. *E. acroporae* Acr-14^T^ genome has been previously made public under accession ID: PJPV00000000

## Results

### 16S rRNA gene phylogeny and *E. acroporae* abundance profiling in different coral species from the Indo-Pacific region

All three strains of *E. acroporae* had only one copy of 16S rRNA gene, compared to seven copies in the *E. montiporae* CL-33^T^ genome (Supplementary Table S3). 16S rRNA gene based phylogeny clustered sequences in a somewhat host-specific way (Supplementary Figure S1). We identified separate clades for *E. montiporae* and *E. acroporae* (Supplementary Figure S1). The closest relative of *E. acroporae* strains was a new species, *Endozoicomonas coralli*, within the same clade, whose genome has not been sequenced yet. Another *Endozoicomonas* species was *Endozoicomonas atrinae* WP70^T^, whose genome has been sequenced.

Relative abundance profiles of *E. acroporae* strains were determined in different coral species from three distinct geographical regions (Fig. 1). We determined that *E. acroporae* strains were abundant in corals from the Red Sea, Saudi Arabia (15-22%); Penghu Archipelago, Taiwan (>50%); and coral reefs in eastern (10-49%) and western (10-75%) Australia. Presence and differential relative abundance of *E. acroporae* strains provide evidence that this species is one of the abundant bacteria among coral holobionts spread across distinct geographic locations.

**Fig. 1.**
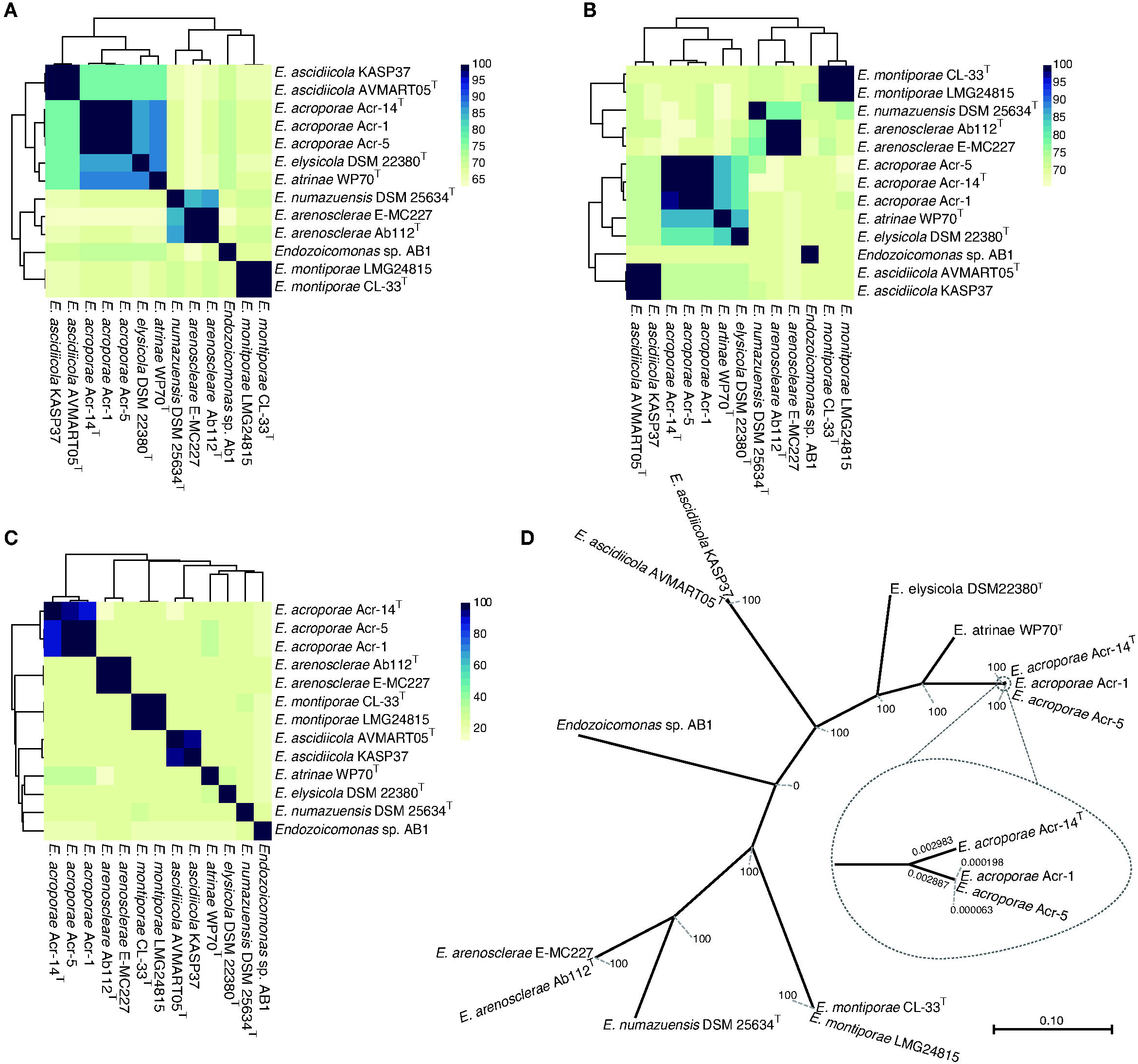
Distribution and abundance profiles of *E. acroporae* strains in the Red Sea, Saudi Arabia; Penghu Archipelago, Taiwan; and eastern and western Australian coral reefs. The green color shows relative abundance of *E. acroporae* strains.

### Genome assembly features

Whole-genome sequencing of *E. acroporae* isolates produced high-quality genome assemblies with completeness >95% and contamination <3% (Supplementary Figure S2). The *E. acroporae* Acr-1 genome was assembled into 299 contigs with a total size of 6,024,033 bp, and we predicted 5,059 coding genes and 77 tRNAs. The *E. acroporae* Acr-5 genome was 6,034,674 bp in length with 295 contigs and 5,101 coding genes and 80 tRNAs. Genome assembly details of *E. acroporae* Acr-14^T^ were 6.048 Mb long, 309 contigs, 5018 coding genes and 81 tRNAs; these have also been reported in an earlier study (Tandon K et al, 2018). A list of all genomes used in this study—along with their genome size (4.049-6.69 Mb) and host (coral, sea slug, comb pen shell and sponge)—is shown in Table S4. All *E. acroporae* genomes (Acr-1, Acr-5 and Acr-14^T^) had similar genome sizes (6.024 −6.048 Mb) and GC contents (49.2, 49.3, and 49.3%, respectively).

### Genomic characteristics of genus *Endozoicomonas*

*Endozoicomonas* species have large genomes ranging from 4.049 Mb (*Endozoicomonas* sp. AB1) to 6.69 Mb long (*E. elysicola* DSM22380). The average genome has ∼5,100 protein coding genes and a GC content of 47.6%. The average DDH, ANI and AAI values among the *Endozoicomonas* species were ∼46, ∼73 and ∼75%, respectively, which are at the lower end of the 62-100% range of interspecies variation within a genus (Kim M et al, 2014), indicating high genomic diversity (Fig. 2A, B, C).

**Fig. 2.**
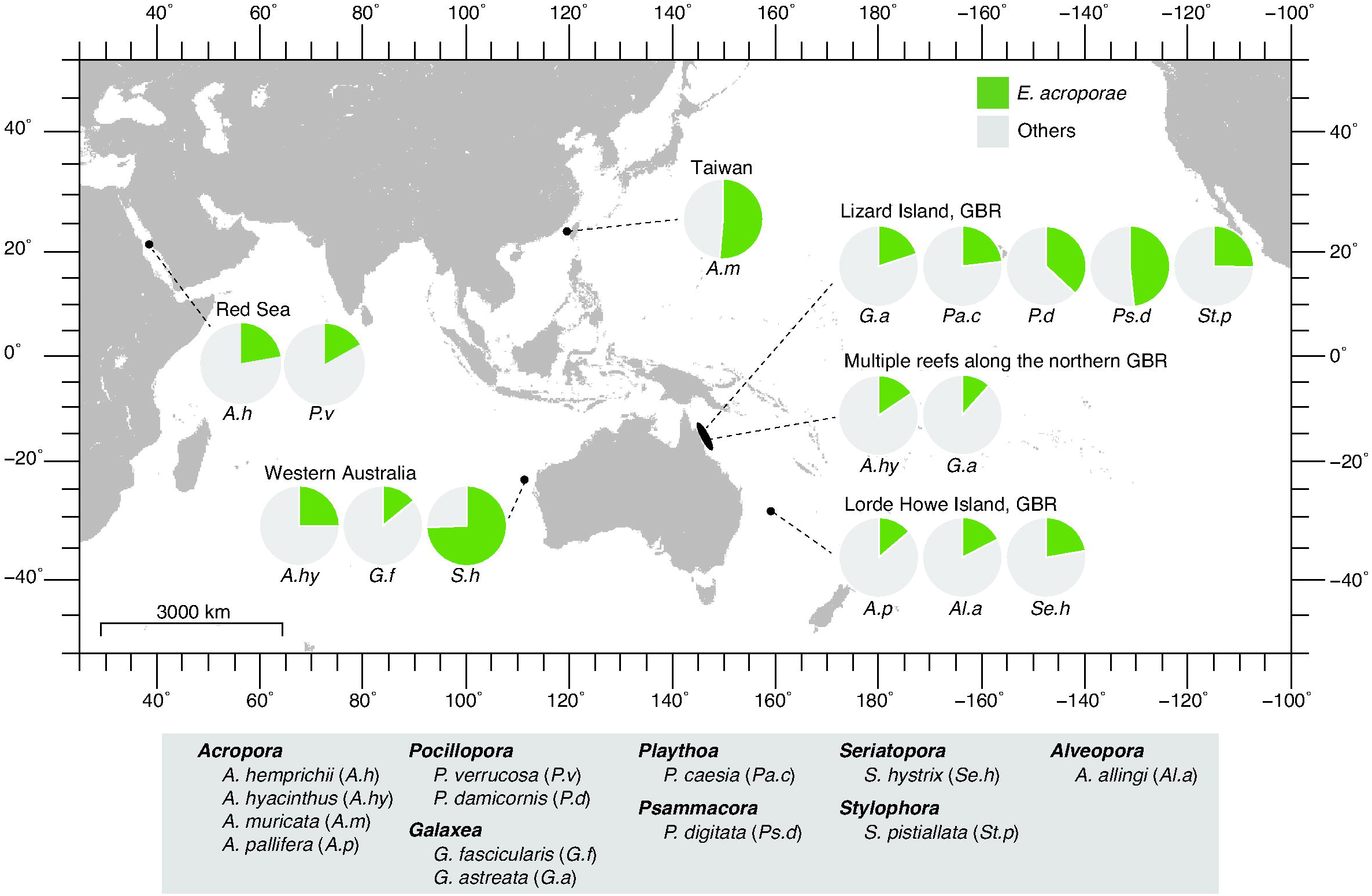
Genomics characteristics of genus *Endozoicomonas.* Heatmaps based on **A)** Average Amino-acid Identity (AAI), **B)** Average Nucleotide Identity (ANI), **C)** DNA-DNA Hybridization (DDH) for all the sequenced genomes from genus *Endozoicomonas.* **D)** Core-genome (308)-based phylogenetic tree.

A diverse array of IS elements were identified when comparing the genomes of *E. acroporae* and *E. montiporae* species. For example, of the total 44 IS elements identified, 22 were only present in *E. acroporae* genomes, while 18 were only present in *E. montiporae* genomes. Relatively few IS elements (ISEc46, ISEcret1, ISEc42 and ISPa18) were present in both species. Each genome also harbored some unique IS elements; among these, Acr-14^T^ had 6 unique IS elements (Supplementary Figure S3).

*E. acroporae* genomes had more genes coding for T3SS with Acr-14^T^ (523), Acr-5(499) and Acr-1 (499) than *E. monitporae* CL-33^T^ (249), *E. atrinae* WP70^T^ (382) or other *Endozoicomonas* species (Supplementary Table S5). All the genomes encode complete pathways for the biogenesis of most amino-acids. We further identified different phage insertions in all *Endozoicomonas* genomes (Supplementary Table S6) and different CRISPR counts in the genomes of *E. acroporae* Acr-14^T^, Acr-5 and Acr-1 (Supplementary Table S3).

### Core genome phylogeny

Core gene (n=308)-based phylogenetic analysis identified a somewhat host-specific phylogeny and also hinted towards a high genomic divergence within the species of genus *Endozoicomonas. Endozoicomonas* genomes isolated from the same host clustered very tightly together, e.g. *E. acroporae* and *E. montiporae* with the coral host and *E. ascidiicola* and *E. arenosclerae* with a sponge host (Fig. 2D). Moreover, *E. numazuensis* and *E. arenosclerae* genomes shared a branch, both being isolated from the sponge. *E. acroporae* Acr-1 and Acr-5 clustered tightly while sharing a branch with *E. acroporae* Acr-14^T^, which confirms that Acr-1 and Acr-5 are closer than Acr-14^T^; the former had ANI value of 99.81%.

### Analyzing RAST subsystems

RAST subsystems; carbohydrates, protein metabolism, and amino acids and derivatives had the highest number of genes, representing an average of 11.69, 10.85 and 13.29% of total annotated genes, respectively (Supplementary Figure S4), in all *Endozoicomonas* genomes. We focused our analysis on two other important subsystems, stress response and sulfur metabolism. The highest number of genes were annotated for oxidative stress (39.16% of stress genes) in the stress response subsystem, followed by unclassified (16%) and detoxification stress (12.53%) (Supplementary Figure S5). Interestingly, in sulfur metabolism, we identified a *dddD* gene homolog capable of metabolizing DMSP into DMS in *E. acroporae* Acr-14^T^, Acr-5, and Acr-1 genomes. This gene was not present in other *Endozoicomonas* genomes (Supplementary Table S4). The *dddD* gene present in *E. acroporae* genomes have two identical CaiB domains (position: 1-416, 439-821) belonging to the coenzyme-A transferase superfamily (Supplementary Fig S7).

### DMSP metabolism operon and links to the central carbon cycle

We identified homologs of the DMSP transcriptional regulator (LysR family), a sulfur transporter belonging to the betaine/carnitine/choline transporter (BCCT), and genes involved in producing acetate from DMSP (Fig. 3A). We discerned a complete pathway with genes arranged in a consecutive manner to form an operon that can yield acetate from DMSP metabolism via three-step enzymatic reactions mediated by DddD, 3-hydroxypropionate dehydrogenase (EC 1.1.1.59), and malonate-semialdehyde dehydrogenase (EC 1.2.1.18) (Fig. 3B) in all the *E. acroporae* genomes with significant gene and protein identity for all the three genes (>97%) and proteins (>97%) (Fig. 3C). Identification of the DMSP cleavage pathway leading to central carbon metabolism suggested that *E. acroporae* species can use DMSP as a carbon source.

**Fig. 3.**
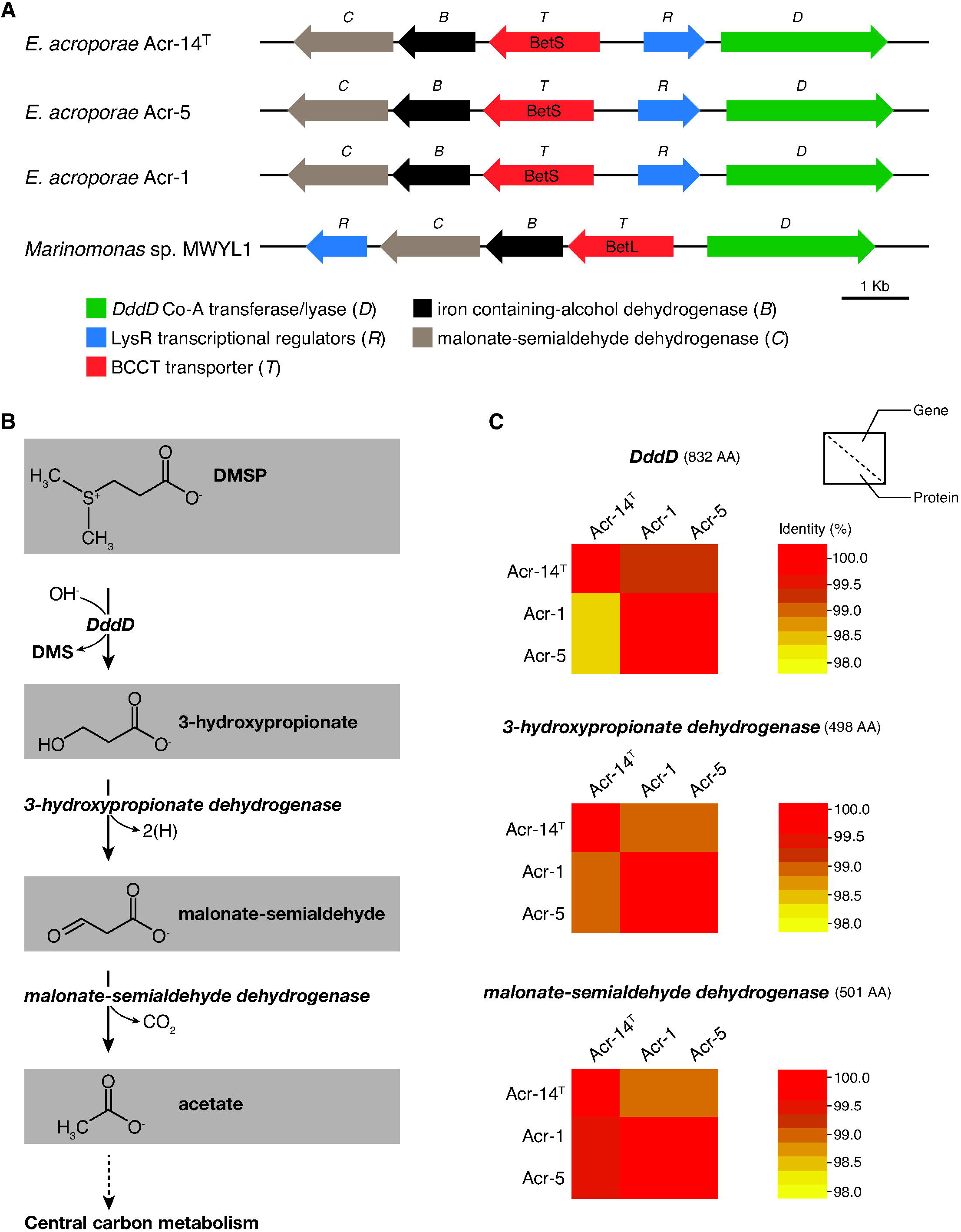
**3A)** DMSP metabolism operon, with genes arranged in an operon manner with in all *E. acroporae* strains, with a similar arrangement in *Marinomonas* sp. MWYL1, with different putative DMSP transporters (*BetS* and *BetL*). **3B)** Metabolic pathway represented by the genes, linking DMSP metabolism to the central carbon cycle, producing acetate from DMSP with the release of DMS. **3C)** Heatmaps based on gene/protein identity for the three enzymes used in the metabolic cycle, DddD, 3-hydroxypropionate dehydrogenase (EC 1.1.1.59), and malonate-semialdehyde dehydrogenase (EC 1.2.1.18).

### *dddD* gene activity and DMS quantification

RT-qPCR was used to examine the expression of the *dddD* gene in *E. acroporae* Acr-14^T^. *dddD* gene expression increased with sampling time in the condition with 1 mM DMSP (Fig. 4A). *dddD* gene expression in this condition was 42.77, 56.52 and 91.37 times higher than samples without DMSP at 16, 24 and 48 hours, respectively (t-test, *p<0.05)*, confirming that *dddD* was active in *E. acroporae*. After confirming the *dddD* gene expression, we quantified the amount of DMS released by *E. acroporae* when incubated in a DMSP-rich environment with a time series detection (0, 24 and 48 hours). We identified that the DMS signal can only be detected in Treatment (a). There was no DMS signal in the control groups (Treatments (b) and (c)) (Fig 4B). A temporal increase in released DMS concentration in Treatment (a) confirmed the ability of *E. acroporae* to metabolize DMSP into DMS.

**Fig. 4.**
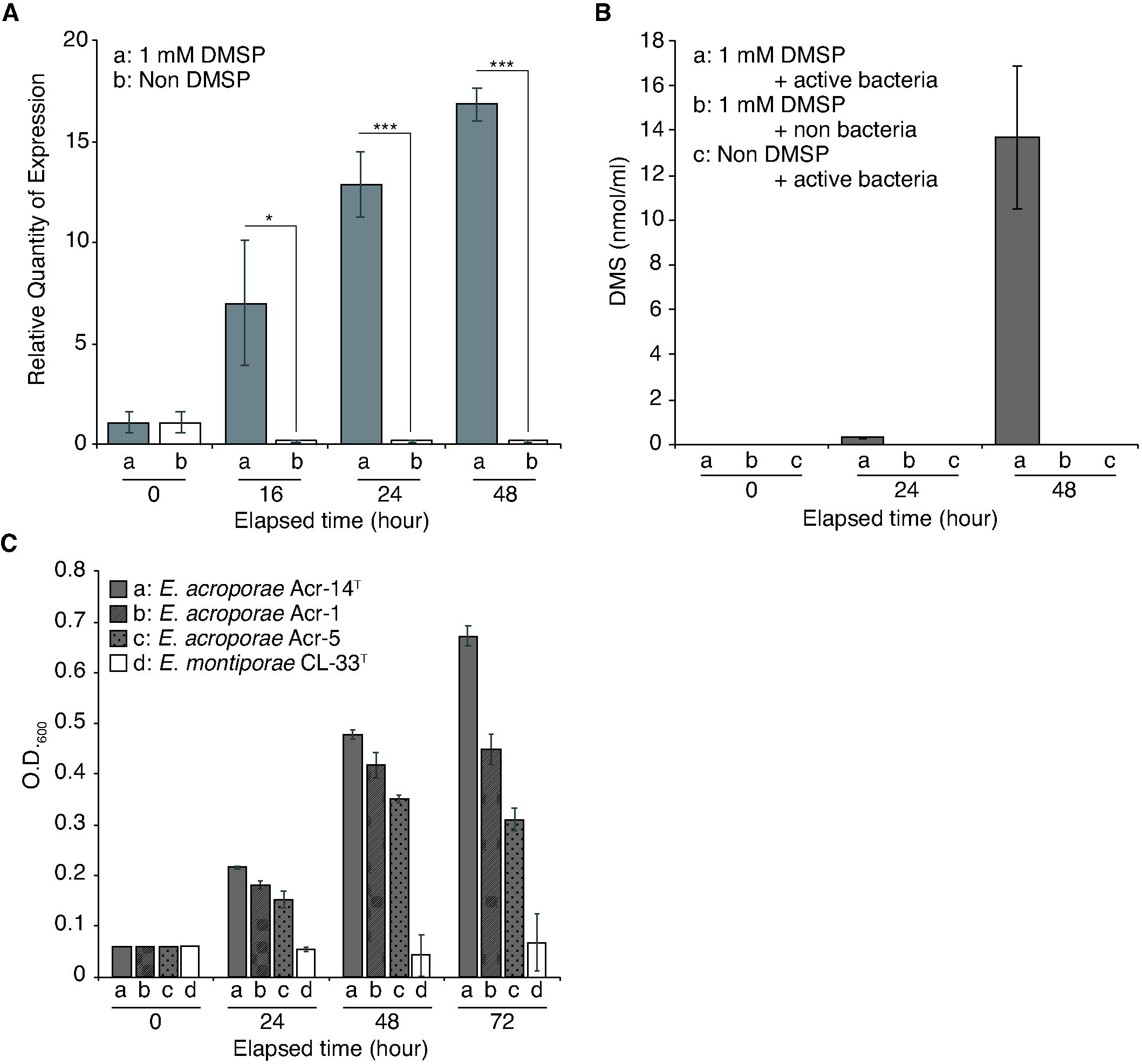
**4A)** RT-qPCR of the *dddD* gene from *E. acroporae* Acr-14^T^ to measure temporal expression change in the presence of 1 mM DMSP. **4B)** GC-based quantification of DMS released from *E. acroporae* Acr-14^T^ cultures in the presence of DMSP (1mM). **4C)** Sole Carbon Test; *E. acroporae* strains can use DMSP as the sole carbon source for their growth and survival, whereas *E. montiporae* CL-33^T^ cannot grow on DMSP as the sole carbon source.

### DMSP as the sole carbon source

*E. acroporae* Acr-14^T^, Acr-1, Acr-5 all showed the ability to use DMSP as the sole carbon source. Acr-14^T^ had a mean OD_600_ value ∼0.7 at the 72-hour time point, which was the highest across all treatments (Fig. 4C). The mean OD_600_ value of Acr-1 and Acr-5 was ∼0.45 and ∼0.3, respectively, after three days of incubation (Fig. 4C). In contrast, *E. montiporae* CL-33^T^ did not show any sign of growth across all time points (Fig. 4C). We than tested whether *E. montiporae* CL-33^T^ could grow under different concentrations of DMSP, equivalent to the same carbon concentration of maltose that can support *E. montiporae* growth (Supplementary Figure S6). The result showed that the *E. montiporae* CL-33^T^ OD_600_ value did not increase across all treatments 0.1, 3, and 7 mM of DMSP after three days of incubation, which further confirmed that *E. montiporae* CL-33^T^ cannot use DMSP as its sole carbon source, but also requires maltose as the carbon source for growth (Supplementary Figure S6).

## Discussion

In this study, we performed genomic (and comparative genomic) and functional assays, and abundance profiling of strains from a newly added species *E. acroporae* in the exclusive marine bacterial genus *Endozoicomonas.* We assembled high quality, nearly complete genomes of *E. acroporae* strains and identified for the first time the *dddD* gene responsible for the previously hypothesized ability of *Endozoicomonas* species to metabolize DMSP. Furthermore, we characterized the functional activity of the identified gene, the ability of *E. acroporae* strains to grow on DMSP as their sole carbon source. This is the first study to establish the role of an *Endozoicomonas* species in the coral sulfur cycle with genomic and functional evidence.

*Endozoicomonas* species have been suggested to play a role in the coral sulfur cycle via DMSP metabolism [8,59]. However, no gene related to DMSP metabolism was found in previously sequenced *Endozoicomonas* genomes (Supplementary Table S4) [30]. Furthermore, using enrichment cultures from coral mucus, tissue and skeleton, certain bacterial species have been reported that can degrade DMSP, including *Spongiobacter (Endozoicomonas) nickelotolerans*, but no gene was identified and it was hypothesized to use a different pathway for DMSP degradation [8]. In this study, we investigated the sulfur metabolism pathway in high quality and nearly complete draft genomes (Supplementary Figure S2) of *E. acroporae* strains and identified a *dddD* gene, which is a Co-A transferase/lyase that cleaves DMSP to volatile DMS [9]. Among the eight identified DMSP lyases in different bacterial species, only *dddD* produces 3-HP and DMS, whereas others form acrylate and DMS [60]. 3-HP is not the usual product of other DMSP lyases, which directly produce acrylate after the cleavage due to their unusual organization of class III Co-A transferase domains [61]. Co-A transferase domains present in the DddD protein are similar to CaiB (Supplementary Figure S7), which is a homodimer capable of adding a Co-A to carnitine [13]. RT-qPCR-based temporal analysis of *E. acroporae* strains in a DMSP-rich environment showed that *dddD* activity increased significantly (Fig. 4A); moreover, the GC analysis also showed that high amounts of DMS were released, providing the evidence that *dddD* is functionally active (Fig. 4B).

Genomic analysis also identified genes arranged in consecutive order to form an operon with DMSP transport, metabolism, and transcriptional regulator genes (Fig. 3A) in all the strains of *E. acroporae. dddT* is a transporter that imports molecules like DMSP [13, 62], and thus has been suggested to be an importer of DMSP. *dddR* is a transcription regulator with the ability to activate the expression of the *dddD* gene in response to DMSP [13]. The functions of *dddB* and *dddC* have not been characterized biochemically, but they are hypothesized to be involved in oxido-reductive function based on their sequences similarity with other oxido-reductases, thereby modifying DMSP either before or after the addition of acyl CoA moiety by *dddD* [13]. Interestingly, the arrangement of genes around *dddD* has been associated to be of the “Pick ‘n Mix” form in different DddD^+^ bacteria, like *Marinomonas sp.* MWYL1, *Sagittus* sp. E37, and *B. cepacia* AMMD [62]; a similar pattern was observed in *E. acroporae* strains as well (Fig. 3A).

The ability of *E. acroporae* strains to use DMSP as their sole carbon source (Fig. 4C) compared to *E. montiporae*—which is unable to utilize DMSP, even at higher concentrations (Supplementary Figure S6)—is further evidence that *E. acroporae* strains can metabolize DMSP and use it for growth and survival. This finding indicates that the DMSP metabolism is directly linked to the central carbon cycle, as we predicted in the genomic analysis (Fig. 3B). Furthermore, it also confirmed the hypothesis that marine bacteria that harbor *dddD* have the ability to grow on DMSP as the sole carbon source [63].

Bacterial genome size can reflect that bacteria’s evolutionary dependency on its host when they are engaged in obligate symbiosis [64]. A striking feature of obligate symbionts is that they have smaller genome sizes than facultative symbionts [65]. However, genomes of *Endozoicomonas* species are relatively long, with a size range of 4.09 Mb to 6.69 Mb; this includes *E. acroporae* strains, which have an average genome length of ∼6.03 Mb, suggesting that they might have free living stage. The possibility that *Endozoicomonas* species have a free living stage is further supported by their large number of coding genes (Avg. ∼5,000 proteins). These features suggest that genome streamlining [66], a notable feature of symbiotic bacteria, does not predominantly occur in genus *Endozoicomonas*. Moreover, a diverse array of phages (Supplementary Table S6) and IS elements (Supplementary Figure S3) provides clues about the infection and colonization histories of different marine hosts, aided by frequent divergence events. Recent studies by Ding JY et al. (2016) [29] reported a high proportion of repeat sequences and also a diverse variety of IS in the genome of *E. montiporae*, which may help the bacterium adapt to the host and also identify the *N-deglycosylation* enzyme that helps penetrate the mucus layer of the host. These findings suggest that *Endozoicomonas* can transition between different symbiotic lifestyles.

We identified a high number of secretory genes (Type III secretion system, T3SS) in the all the genomes of *E. acroporae* (Supplementary Table S5) that may help transport organic maromolecules between the host and symbiont. The T3SS system helps bacteria interact with their host, and a complete gene set for the assembly of this vital secretory system was observed in all the genomes used in this study. *E. acroporae* has secretory genes that regulate host mechanisms, similar to those of *E. montiporae*, and can increase the chances that bacteria survive in the host while improving the host fitness. Additionally, other genes found in *E. acroporae* may also provide clues about the survival strategy of *E. acroporae* in hosts. For example, identification of a catalase gene in *E. acroporae* strains—along with phosphoenolpyruvate synthase and 7,8-dihydro-8-oxoguanine triphosphatase as secretory genes from T3SS—can help the bacterium survive by scavenging H_2_O_2_, modulating the gluconeogeneis and confer resistance from oxidative stresses to the host respectively. Up-regulating genes involved in gluconeogenesis have been proposed as a response to stress-induced starvation in corals [67].

Phylogenetic analysis based on a core-genome (308 genes) clustered the species in a somewhat host-specific manner, as it has done in previous studies [30, 31]. However, it was interesting to see that, even when the host type is the same (i.e. stony coral), *E. montiporae* and *E. acroporae* did not share any branch, and their strains cluster tightly within their clade, which suggests that the co-diversification between host and symbiont is complex in nature (Fig. 2D). Moreover, a highly reduced core genome along with similar ANI, DDH and AAI at the lower end of inter species range give clues about the interspecies diversity within genus *Endozoicomonas* (Fig. 2A, B, C).

With the high abundance of *Endozoicomonas* species reported in coral holobionts across the globe and very few culturable isolates known to date, it has been hypothesized that *Endozoicomonas* is either species specific or can have a large host range. In our study, we identified as being *E. acroporae* prevalent in diverse coral species from geographically distant locations in the Indo-Pacific region. Relative abundance varies among coral species (Fig. 1). The presence of *E. acroporae* in different coral species supports the idea that this new species has a large host range in the genus *Endozoicomonas;* it also highlights the importance of studying this genus in greater detail. Identification of other isolates that can be cultured will enhance our understanding of this diverse marine genus. This further points to the need to advance our culturing techniques and combine the sequencing approaches with culturomics.

*Endozoicomonas* species have only been isolated from marine invertebrates; hence it is reasonable to believe that *Endozoicomonas* and their hosts experiences ever-present oxidative stress [69] in the marine environment. Identification of a high percentage of oxidative stress response genes in all the genomes of *Endozoicomonas* provides clues for understanding its adaptation to the marine environment and potential ability to mitigate oxidative stress. Moreover, earlier studies have reported that DMSP, which has antioxidant properties, and its breakdown products are functionally important in the coral stress response [69]. In our study, we also identified 7,8-dihydro-8-oxoguanine triphosphatase as a T3SS effector protein, which might provide resistance against mitochondrial dysfunction and spontaneous mutagenesis [29, 70] to the coral host during thermal stress, which is also an indication of a putative symbiotic association between E. *acroporae* and *A. muricata.*

## Conclusion

*Endozoicomonas* species have been long hypothesized to play a role in the coral sulfur cycle, and this study provides genomic and functional evidence for the first time to support this hypothesis. *E. acroporae* strains can not only metabolize DMSP to produce DMS, but also use it as the sole carbon source for growth and survival. In our study we also identified the first DMSP-related operon in *E. acroporae*, which links DMSP metabolism to the central carbon cycle. Furthermore, the presence of stress responsive genes at higher proportions gives clues about how this genus adapted in marine environments. Since very few species in this diverse marine genus have been cultured to date, we believe that there are still many other functional aspects, and more focus on this genus in regards to coral reef health can provide better insights in the near future.

## Acknowledgements

This study was supported by funding from Academia Sinica. K.T would like to acknowledge the Taiwan International Graduate Program (TIGP) for its fellowship towards his graduate studies. The authors would like to acknowledge support from Dr. Shu-Fen Chiou during the GC experiment. We would like to thank Noah Last of Third Draft Editing for his English language editing.

## Author contribution

K.T and S.L.T conceived the idea for this study. K.T assembled the genomes, performed the bioinformatics analysis, and wrote the manuscript. P.W.C cultured the strains and performed the RT-qPCR analysis. C.Y.L and Y.F.C performed GC experiment and analysis. C.Y.L performed the sole carbon test and analysis. N.W and S.Y helped write the manuscript and modify the illustrations. P.Y.C, H.Y.C, and M.S.C helped with the GC experiments and provided the instruments for conducting the experiment. W.M.C provided the cultures. S.L.T supervised the overall study. All authors read and approved the manuscript.

