## Supplementary data for "Comparative genomics: Dominant coral-bacterium *Endozoicomonas acroporae* metabolizes dimethylsulfoniopropionate (DMSP)"

### Supplementary Figures and Tables Legends

Supplementary Figure S1. 16S rRNA gene-based phylogenetic tree for all 16S rRNA sequences (80 in number) from genus *Endozoicomonas* whose source of isolation is known. *E. acroporae* strains belong to the clade coral group 1 *Acropora muicata,* along with a newly discovered species *Endozoicomonas coralli* strain Acr-12.

Supplementary Figure S2. CheckM plots for *Endozoicomonas acroprae* strains Acr-1, Acr-5 and Acr-14$^T$.

Supplementary Figure S3. Distribution of different IS elements in *E. acroporae* and *E. montiporae* strains. Color codes represent the different combination and numbers represent the copies of each IS element identified in the genome.

Supplementary Figure S4. Proportion of different subsystem annotations by RAST Server on all the genomes used in this study.

Supplementary Figure S5. Proportion of different stress responsive genes annotated by RAST in the Stress Response Subsystem. Oxidative stress response genes account for ~40% of the total genes annotated.

Supplementary Figure S6. $OD_{600}$ values as an indicator of *E. montiporae* CL-33$^T$ growth with different concentrations of DMSP (d, e, and f), maltose (g) and *E. acroporae* Acr-14$^T$ with 0.1 mM of DMSP. *E. montiporae* only grew with maltose as the carbon source, and *E. acroporae* grew well with DMSP.

Supplementary Figure S7. Representation of functional domains specific hit (CaiB) and superfamily (CoA_transf_3) present in DddD protein of *E. acroporae* strains.

Supplementary Table S1. Composition of Modified Marine Broth, Version 4 (MMBV4*).

Supplementary Table S2. Composition of minimal medium for *Endozoicomonas*.

Supplementary Table S3. Genome assembly characteristics of *E. acroporae* strains.

Supplementary Table S4. List of genomes used in this study for comparative genomic analysis with their genome size, host and presence of *dddD* gene.

Supplementary Table S5. Count of Type III secretion system (T3SS) genes identified in genomes of *Endozoicomonas* species.

Supplementary Table S6. Phage insertions identified in genomes of genus *Endozoicomonas*. only intact phages are annotated.
