## Supplementary figures and images for "Comparative genomics: Dominant coral-bacterium *Endozoicomonas acroporae* metabolizes dimethylsulfoniopropionate (DMSP)"

### Supplementary Figure S1

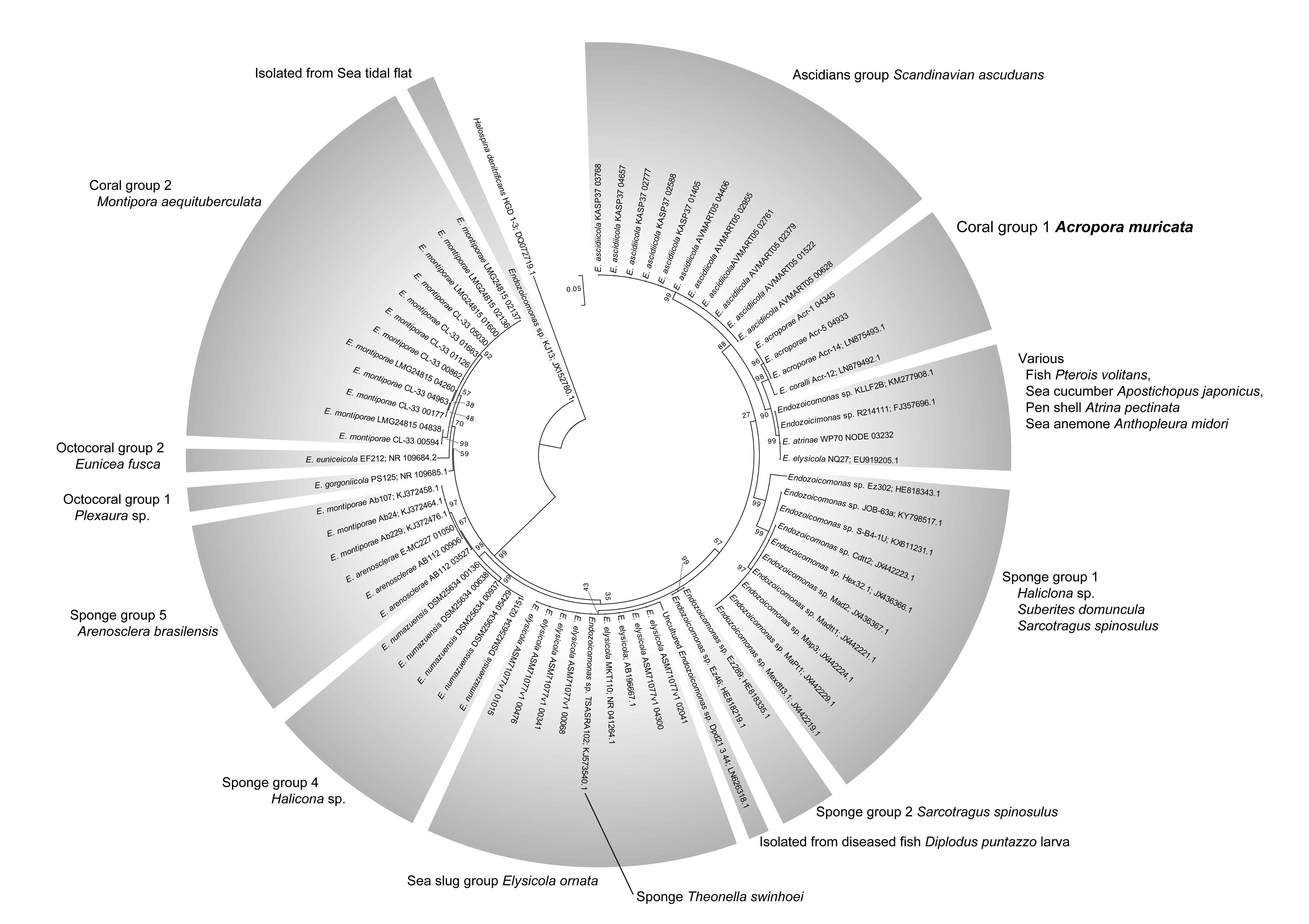

### Supplementary Figure S2

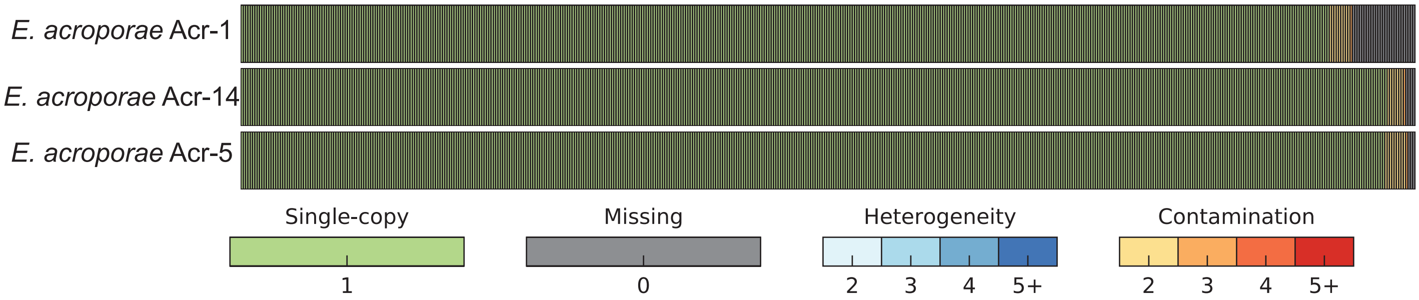

### Supplementary Figure S4

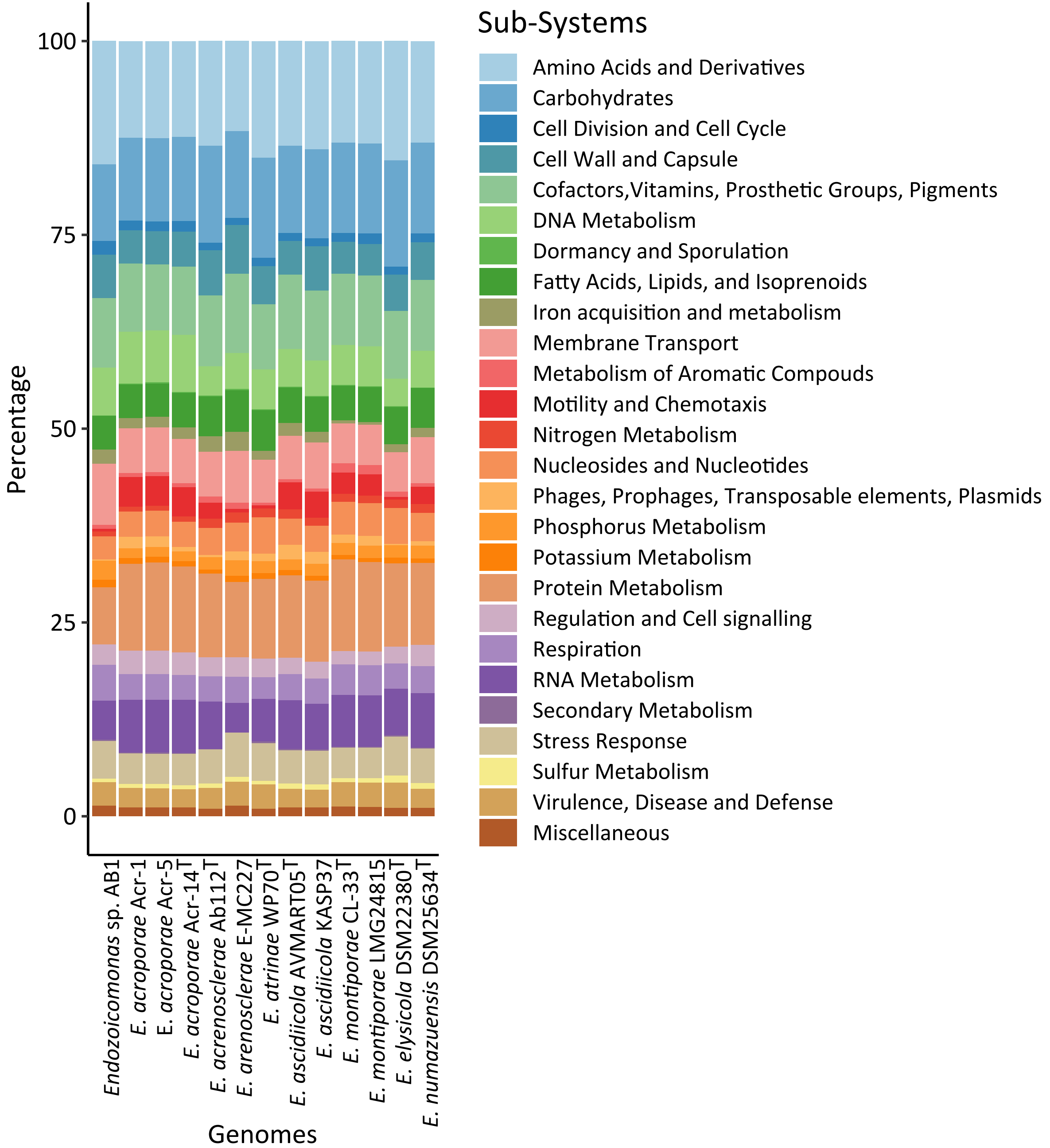

### Supplementary Figure S5

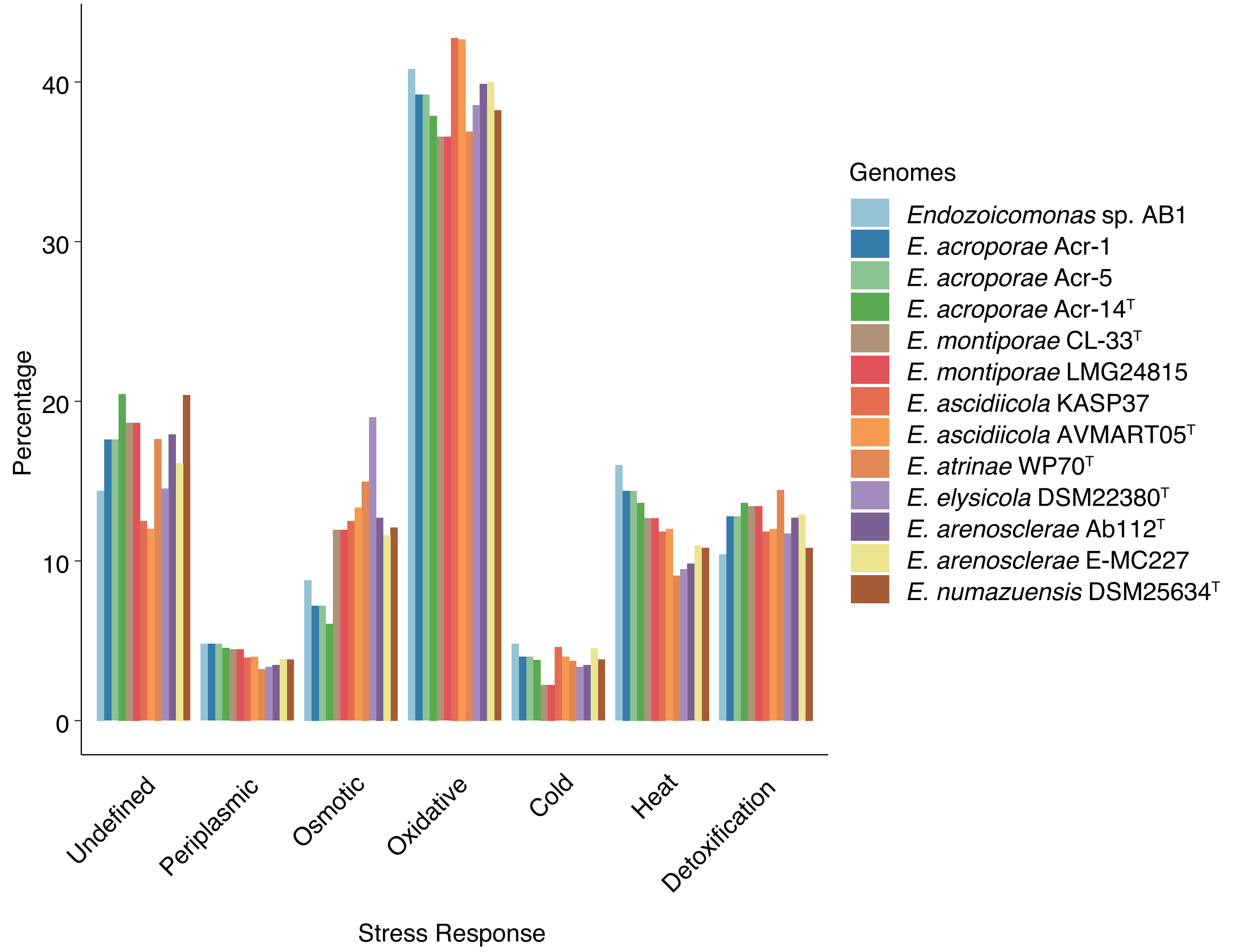

### Supplementary Figure S6

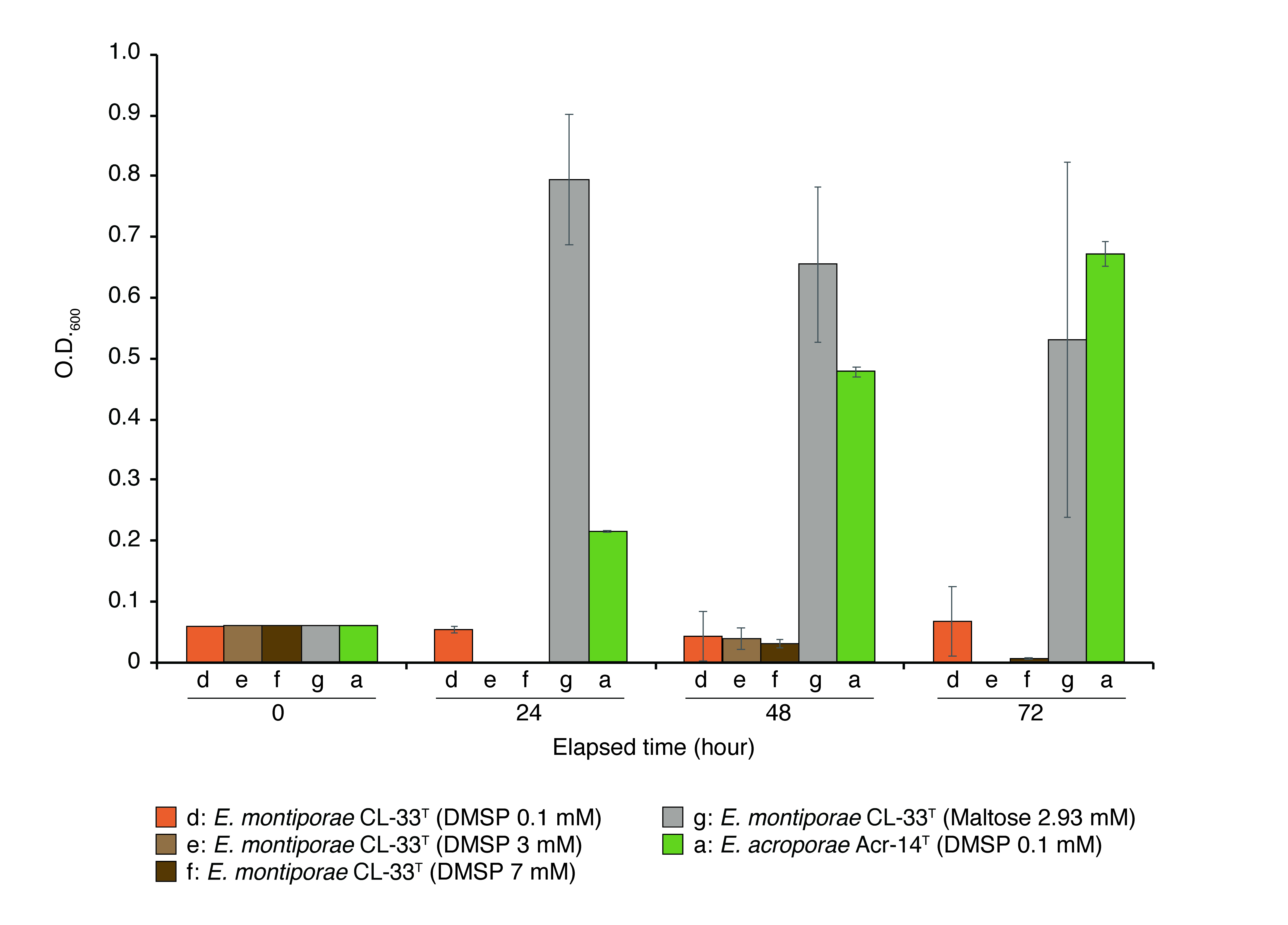

### Supplementary Figure S7

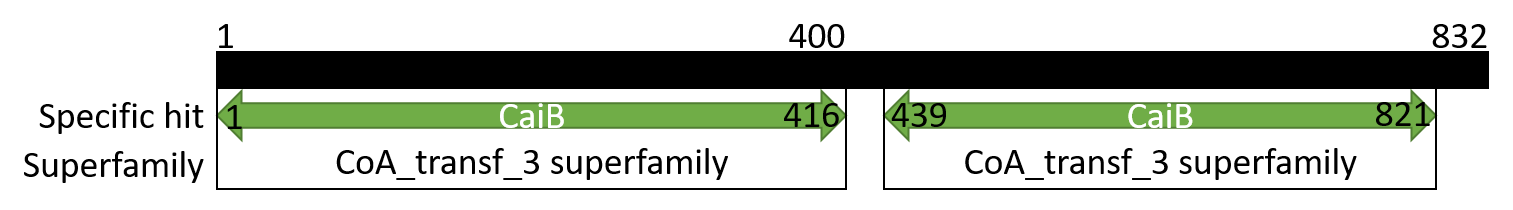
