## Supplementary Figure S3 for "Comparative genomics: Dominant coral-bacterium *Endozoicomonas acroporae* metabolizes dimethylsulfoniopropionate (DMSP)"

| IS element | ISEc46 | ISEcret1 | ISEc42 | ISPa18 | IS1618 | ISAbe14 | ISAeme8 | ISAeme9 | ISCARN97 | ISEcret6 | ISEnu2 | ISKpn25 | ISRop1 | ISXne4 | ISPa38 | ISPsy41 | ISShfr7 | ISShwo5 | ISCARN4 | ISEcret9 | ISNtsp2 | ISPin5 | ISPlu2 | ISPpr13 | ISShes1 | ISSod15 | ISSpu1 | ISSpu5 | ISThma1 | ISPlu19 | ISSm4 | ISBps1 | ISCARN64 | ISGes1 | ISGur6 | ISHbo2 | ISHhu3 | ISPa82 | ISPsy44 | ISEcret2 | ISEc18 | ISEc40 | ISPlu3 | ISVa15 |
|---|---|---|---|---|---|---|---|---|---|---|---|---|---|---|---|---|---|---|---|---|---|---|---|---|---|---|---|---|---|---|---|---|---|---|---|---|---|---|---|---|---|---|---|---|
| Acr-1 | 1 | 2 | | 1 | 2 | 1 | 4 | 3 | 1 | 3 | 2 | 1 | 1 | 2 | 1 | 1 | 1 | 1 | | | | | | | | | | | | | | 1 | 1 | | | | | | | | | | | |
| Acr-5 | 1 | 2 | | 1 | 2 | 1 | 4 | 3 | 1 | 3 | 1 | 2 | 1 | 1 | 1 | 1 | 1 | 1 | | | | | | | | | | | | | | | | | | | | | | | | | | |
| Acr-14$^T$ | 1 | 3 | 1 | 3 | 1 | 1 | 2 | 9 | 2 | 2 | 2 | 2 | 1 | 5 | | | | | | | | | | | | | | | | | | | | 1 | 5 | 1 | 1 | 1 | 1 | | | | | |
| CL-33$^T$ | 2 | 1 | 1 | | | | | | | | | | | | | | | | 3 | 4 | 5 | 3 | 3 | 7 | 1 | 1 | 1 | 3 | 5 | 5 | 1 | | | | | | | | | 1 | | | | |
| LMG-24815 | 2 | 1 | 1 | 1 | | | | | | | | | | | | | | | 4 | 3 | 3 | 2 | 2 | 5 | 1 | 1 | 1 | 3 | 3 | 3 | 1 | | | | | | | | | | 1 | 4 | 1 | 1 |

Shared in all

Between *E. acroporae* and *E. montiporae*

Shared in *E. acroporae*

Only shared in *E. acroporae* Acr-1 and Acr-5

Only shared in *E. montiporae*

Unique in a strain
