## Supplementary Table S1 for "Comparative genomics: Dominant coral-bacterium *Endozoicomonas acroporae* metabolizes dimethylsulfoniopropionate (DMSP)"

Supplementary Table S1. Composition of Modified Marine Broth, Version 4 (MMBV4*)*.*

| **Ingredients** | **Volume (concentration)** |
| --- | --- |
| HEPES | 5.95 g/l (25mM) |
| NaCl | 19.45 g/l (0.33M) |
| MgCl_2_∙6H_2_O | 18.79 g/l (92.4mM) |
| Na_2_SO_4_ | 3.24 g/l (22.8mM) |
| KCl | 0.55 g/l (7.377mM) |
| CaCl_2_ | 0.12 g/l (1.08mM) |
| NaHCO_3_ | 0.16 g/l (1.9mM) |
| Peptone | 5 g/l |
| Yeast extract | 1 g/l |
| Trace metal solution | 1 ml/l |
| NP cocktail stock** | 1 ml/l |
| pH | 7.2 |

*MMBV4 was modified from Ding JY et al., 2016.

**NP cocktail stock contained KNO_3_(89.21g/L), NH_4_Cl(47.2g/L) and NaH_2_PO_4_∙H_2_O(4.35g/L).
