## Supplementary Table S2 for "Comparative genomics: Dominant coral-bacterium *Endozoicomonas acroporae* metabolizes dimethylsulfoniopropionate (DMSP)"

Supplementary Table S2. Composition of minimal medium for *Endozoicomonas*.

| **Ingredients** | **Volume (concentration)** |
| --- | --- |
| 5x ASW stock | 200 ml |
| CaCl_2_ | 1g/l (9 mM) |
| NaHCO_3_ | 168 mg/l (2 mM) |
| HEPES | 2.38 g/l (10 mM) |
| NO_3_-Pi stock | 1 ml |
| Trace metal solution | 1 ml |
| Wolf’s vitamin solution | 1 ml |
| Sodium alginate | 4g/l (23.64 mM) |
| Asparagine | 132 mg/l (1mM) |
| Histidine | 223 mg/l (1.44mM) |
| pH | 7.4 |
