## Supplementary Table S3 for "Comparative genomics: Dominant coral-bacterium *Endozoicomonas acroporae* metabolizes dimethylsulfoniopropionate (DMSP)"

Supplementary Table S3. Genome assembly characteristics of *E. acroporae* strains.

| **Genome Characteristics** | ***E. acoporae*  Acr-14^T^** | ***E. acroporae*  Acr-5** | ***E. acroporae* Acr-1** |
| --- | --- | --- | --- |
| Size (Mbp) | 6.048 | 6.034 | 6.024 |
| GC content | 49.16% | 49.3% | 49.2% |
| N50 | 47,658(bp) | 52,448(bp) | 56,565(bp) |
| No. of rRNA | 5 (16S x 1, 5S x 4) | 7 (16S x 1, 5S x 6) | 6 (16S x 1, 5S x 5) |
| No. of tRNA | 81 | 80 | 77 |
| No. of Genes | 5,104 | 5,190 | 5,144 |
| No. of CDS | 5,018 | 5,101 | 5,059 |
| CRISPRs | 4 | 3 | 2 |
