## Supplementary Table S4 for "Comparative genomics: Dominant coral-bacterium *Endozoicomonas acroporae* metabolizes dimethylsulfoniopropionate (DMSP)"

Supplementary Table S4. List of genomes used in this study for comparative genomic analysis with their genome size, host and presence of *dddD* gene.

| **Genomes** | **Genome Size**  **(# contigs)** | **Host** | ***dddD* gene** |
| --- | --- | --- | --- |
| *Endozoicomonas acroporae* Acr-1 (this study) | 6.024 Mb (299) | Coral *(Acropora muricata)* | **Present** |
| *Endozoicomonas acroporae* Acr-5 (this study) | 6.034 Mb (295) | Coral (*Acropora muricata)* | **Present** |
| *Endozoicomonas acroporae* Acr-14^T^(this study) | 6.049 Mb (309) | Coral *(Acropora muricata)* | **Present** |
| *Endozoicomonas montiporae* CL-33^T^ | 5.430 Mb (1) | Coral (*Montipora aequituberculata)* | Absent |
| *Endozoicomonas montiporae* LMG24815 | 5.602 Mb (20) | Coral (*Montipora aequituberculata)* | Absent |
| *Endozoicomonas atrinae* WP70 ^T^ | 6.690 Mb (985) | Comb pen shell (*Atrina pectinate*) | Absent |
| *Endozoicomonas elysicola* DSM22380 ^T^ | 5.606 Mb (2) | Sea Slug (*Elysia ornate)* | Absent |
| *Endozoicomonas* sp. AB1 | 4.049 Mb (272) | *Bugula neritina* AB1 | Absent |
| *Endozoicomonas ascidiicola* AVMART05 ^T^ | 6.135 Mb (36) | Ascidians (Tunicata, Ascidiaceae) | Absent |
| *Endozoicomonas ascidiicola* KASP37 | 6.512 Mb (34) | Ascidians (Tunicata, Ascidiaceae) | Absent |
| *Endozoicomonas numazuensis* DSM25634 ^T^ | 6.340 Mb (31) | Marine sponge | Absent |
| *Endozoicomonas arenosclerae* Ab112 ^T^ | 6.453 Mb (328) | Marine sponge (*Arenosclera brasiliensis)* | Absent |
| *Endozoicomonas arenosclerae* E-MC227 | 6.216 (2501) | Marine sponge (*Arenosclera brasiliensis)* | Absent |
