## Supplementary Table S5 for "Comparative genomics: Dominant coral-bacterium *Endozoicomonas acroporae* metabolizes dimethylsulfoniopropionate (DMSP)"

Supplementary Table S5. Count of Type III secretion system (T3SS) genes identified in genomes of *Endozoicomonas* species.

| **Genomes** | **T3SS gene count** |
| --- | --- |
| *Endozoicomonas acroporae* Acr-1 (this study) | 499 |
| *Endozoicomonas acroporae* Acr-5 (this study) | 499 |
| *Endozoicomonas acroporae* Acr-14^T^( this study) | 523 |
| *Endozoicomonas montiporae* CL-33^T^ | 249 |
| *Endozoicomonas montiporae* LMG24815 | 258 |
| *Endozoicomonas atrinae* WP70 ^T^ | 381 |
| *Endozoicomonas elysicola* DSM22380 ^T^ | 314 |
| *Endozoicomonas* sp. AB1 | 165 |
| *Endozoicomonas ascidiicola* AVMART05 ^T^ | 343 |
| *Endozoicomonas ascidiicola* KASP37 | 360 |
| *Endozoicomonas numazuensis* DSM25634 ^T^ | 301 |
| *Endozoicomonas arenosclerae* Ab112 ^T^ | 297 |
| *Endozoicomonas arenosclerae* E-MC227 | 309 |
