## Supplementary Table S6 for "Comparative genomics: Dominant coral-bacterium *Endozoicomonas acroporae* metabolizes dimethylsulfoniopropionate (DMSP)"

Supplementary Table S6. Phage insertions identified in genomes of genus *Endozoicomonas*. only intact phages are annotated.

| **Genome** | **Intact** | **Incomplete** | **Questionable** | **Phage Annotation (Intact only)** |
| --- | --- | --- | --- | --- |
| *E. acroporae* Acr-14^T^ | 2 | 2 | 1 | PHAGE_Pseudo_MD8_NC_031091, PHAGE_Paenib_Tripp_NC_028930 |
| *E. acroporae* Acr-5 | 3 | 3 | 1 | PHAGE_Bacill_SP_15_NC_031245, PHAGE_Salmon_phSE_2_NC_031026, PHAGE_Entero_Arya_NC_03148 |
| *E.* *acroporae* Acr-1 | 4 | 1 | 2 | PHAGE_Enter_Arya_NC031048, PHAGE_Propio_PFR2_NC_031108, PHAGE_Entero_UAB_Phi20_NC_031019, PHAGE_Stx2_vB_EcoP_NC_027984 |
| **Other Genomes** |  |  |  |  |
| *E.* sp. AB1-5 | 0 | 4 | 1 |  |
| *E. arenosclerae* Ab112 ^T^ | 1 | 2 | 2 | PHAGE_Clostr_c_st_NC_007581 |
| *E. areonsclerae* E-MC227 | 1 | 3 | 1 | PHAGE_Entero_Phi27_NC_003356 |
| *E. elysicola* DSM22380 ^T^ | 0 | 1 | 0 |  |
| *E. montiporae* CL-33^T^ | 2 | 4 | 1 | PHAGE_Clostr_phiCT453B_NC_029004, PHAGE_Salmon_SJ46_NC_031129 |
| *E. montiporae* LMG 24815 | 2 | 4 | 3 | PHAGE_Entero_Arya_NC_031048 (2) |
| *E. ascidiicola* AVMART05 ^T^ | 3 | 2 | 1 | PHAGE_Mesorh_phagevB_MloP_Lo5R7ANS_NC_02543, PHAGE_Escher_vB_ECO1230_10_NC_027995, PHAGE_Pseudo_NP1_NC_031058 |
| *E. ascidiicola* KASP37 | 0 | 5 | 2 |  |
| *E. atrinae* WP70 ^T^ | 0 | 16 | 3 |  |
| *E. numazuensis* DSM25634 ^T^ | 1 | 1 | 1 | PHAGE_Escher_D108_NC_013594 |
